# Characterising the Modulatory Role of Ikebana in Flower-Dependent Cell Competition

**DOI:** 10.1101/2024.04.13.589362

**Authors:** Mariana Marques-Reis, Andrés Gutiérrez-García, Carlos Flores Clara, Irene Argudo, Matthias Stefan Eggel, Barbara Hauert, Catarina Brás-Pereira, Eduardo Moreno

## Abstract

Tissues encompass a quality control mechanism that promotes their optimal state. This mechanism, designated cell competition, is characterised by the elimination of suboptimal yet viable cells when they are near healthier cells within the same tissue compartment.

This study explores Flower-dependent cell competition and introduces Ikebana as a novel player. The differential expression of the *flower* isoforms labels cells as winners or losers, influencing their fate in diverse contexts, including eye development, traumatic brain injury, and Alzheimer’s disease. Ikebana, ubiquitously produced in wing imaginal discs and adult brains, modulates loser cell elimination. Reduction of *ikebana* expression correlates with an increased number of loser cells, while its overexpression in the Alzheimer’s disease model reduces the number of Flower LoseB-positive cells.

We suggest that Ikebana protects loser cell elimination, particularly when excessive elimination of loser cells can compromise tissue function. Thus, Ikebana might be a potential therapeutic target for modulating Flower LoseB expression.

## Introduction

The Darwinian principle of “survival of the fittest” has transcended the animal kingdom to encompass the cellular landscape within their bodies (Moreno and Rhiner 2014). Current knowledge recognises that cells within the same tissue compartment engage in a phenomenon known as Cell Competition, a process elucidated as early as 1975 with the observation of slow-proliferating *minute* ribosomal mutants being outcompeted by normally proliferating cells (Morata and Ripoll 1975). This phenomenon extends beyond mere cell proliferation differences (Böhni et al. 1999), leading to a broader definition of cell competition— the elimination of viable yet suboptimal cells when juxtaposed with fitter counterparts within the same tissue compartment.

Cell competition has been documented across various tissues in organisms ranging from *Drosophila* and mice to zebrafish and humans, with implications in neurodegenerative diseases and cancer biology (Akieda et al. 2019; Coelho et al. 2018; Eisenhoffer et al. 2012; Madan et al. 2019; Oliver et al. 2004; Villa del Campo et al. 2014; Walderich et al. 2016). The prevailing consensus in the field identifies three major mechanisms of cell competition. First is competition for survival factors, exemplified by decapentaplegic capture during wing imaginal disc development, where cells closer to essential factors, or more able to capture them, gain a competitive advantage (Moreno, Basler, and Morata 2002). Second, mechanical cell competition, where cells compete for space, and the resistance to mechanical forces determines their fate, as seen in the elimination of cells converging to the midline in the *Drosophila* thorax (Brás-Pereira and Moreno 2018; Levayer, Dupont, and Moreno 2016; Moreno et al. 2019). Lastly, cells may compete based on their fitness state, influenced by factors such as ageing (Merino et al. 2015) or exposure to toxic environments (Coelho et al. 2018). This leads to the elimination of suboptimal cells (losers) and their potential replacement by fitter cells when in the vicinity of healthier (winner) cells. Fitness fingerprints are proteins that recognise and define fitness states and are thought to play crucial roles in the execution of cell competition. The fitness fingerprints that have been identified to date include Flower (Rhiner et al. 2010), Slit-Robo-Ena (Vaughen and Igaki 2016), Spätzle-Toll (Alpar, Bergantiños, and Johnston 2018; Germani et al. 2018; Tan et al. 2008), Sas-PTP10D (Yamamoto et al. 2017), and Flamingo (Bosch, Cho, and Axelrod n.d.).

This study delves into Flower-dependent cell competition, which relies on the transmembrane protein Flower (Rhiner et al. 2010). Although Flower has been initially described as a calcium channel (Yao et al. 2009, 2017), such function appears irrelevant to cell competition (Coelho and Moreno 2020; Madan et al. 2019). In *Drosophila*, Flower exists in three isoforms – Flower LoseA and Flower LoseB, marking loser cells, and Flower Ubi, marking winner cells (Rhiner et al. 2010). The differential expression of these isoforms determines the fate of a cell as a winner or loser. The expression of *flower lose* isoforms in *Drosophila* causes the elimination of suboptimal cells in various contexts, including during eye development (Merino et al. 2013), in response to traumatic brain injury (Moreno et al. 2015), and in an Alzheimer’s disease model (Coelho et al. 2018; Coelho and Moreno 2019, 2020). The significance of Flower extends beyond *Drosophila*, with orthologues described in mice and humans (Madan et al. 2019; Petrova et al. 2012).

In *Drosophila*, loser cells can experience various fates: if they express *SPARC*, they are protected from elimination; on the other hand, if they express the fitness checkpoint *azot*, they are then marked to die (Merino et al. 2015; Portela et al. 2010). Azot, a predicted calcium-calmodulin, activates *hid* and the subsequent machinery, leading to apoptosis (Merino et al. 2015). However, the mechanisms underlying neighbour recognition as winners or losers, the criteria for selecting loser cells for elimination, and the interplay between Flower, Azot, and apoptosis remain areas of ongoing exploration.

This study introduces Ikebana as a new participant in the landscape of cell competition, which acts as a key regulator in determining cell fate. In scenarios where competition is intensified, as seen in Alzheimer’s disease, increasing *ikebana* expression reduces the number of cells marked as losers. Our findings position Ikebana as a modulator of loser cell elimination, which might be particularly relevant in contexts where excessive labelling of cells as losers could compromise tissue function. With predicted human orthologs, Ikebana presents a potential avenue for developing novel therapies to modulate Flower expression and cell competition.

## Results

### Ikebana is predicted to interact with Flower and is basally produced in the wing-imaginal discs and adult brain

Our understanding of the Flower-dependent cell competition pathway remains limited, so our objective was to characterise novel participants in this process. Based on a co-affinity precipitation assay coupled with mass spectrometry, we focused on proteins predicted to interact with Flower, which revealed 34 proteins as potential interactors (Guruharsha et al. 2011). From this list, only Mec2 and CG15098 were expected to be transmembrane proteins, which is crucial as communication and fitness status labelling likely occur at the plasma membrane. Out of these two proteins, we previously found that only *CG15098* is upregulated in loser cells in a microarray analysis designed to identify genes differentially expressed during cell competition (Rhiner et al. 2010). We have named this gene *ikebana*, drawing inspiration from the Japanese art of flower arrangements. Ikebana has four transmembrane domains, with the -N and -C terminus predicted to be intracellular (*Figure* 1A), and features two predicted domains—Mtp and DUF4728—placing it within the family of tetraspanin-pasiflora proteins (Deligiannaki et al. 2015).

**Figure 1.**
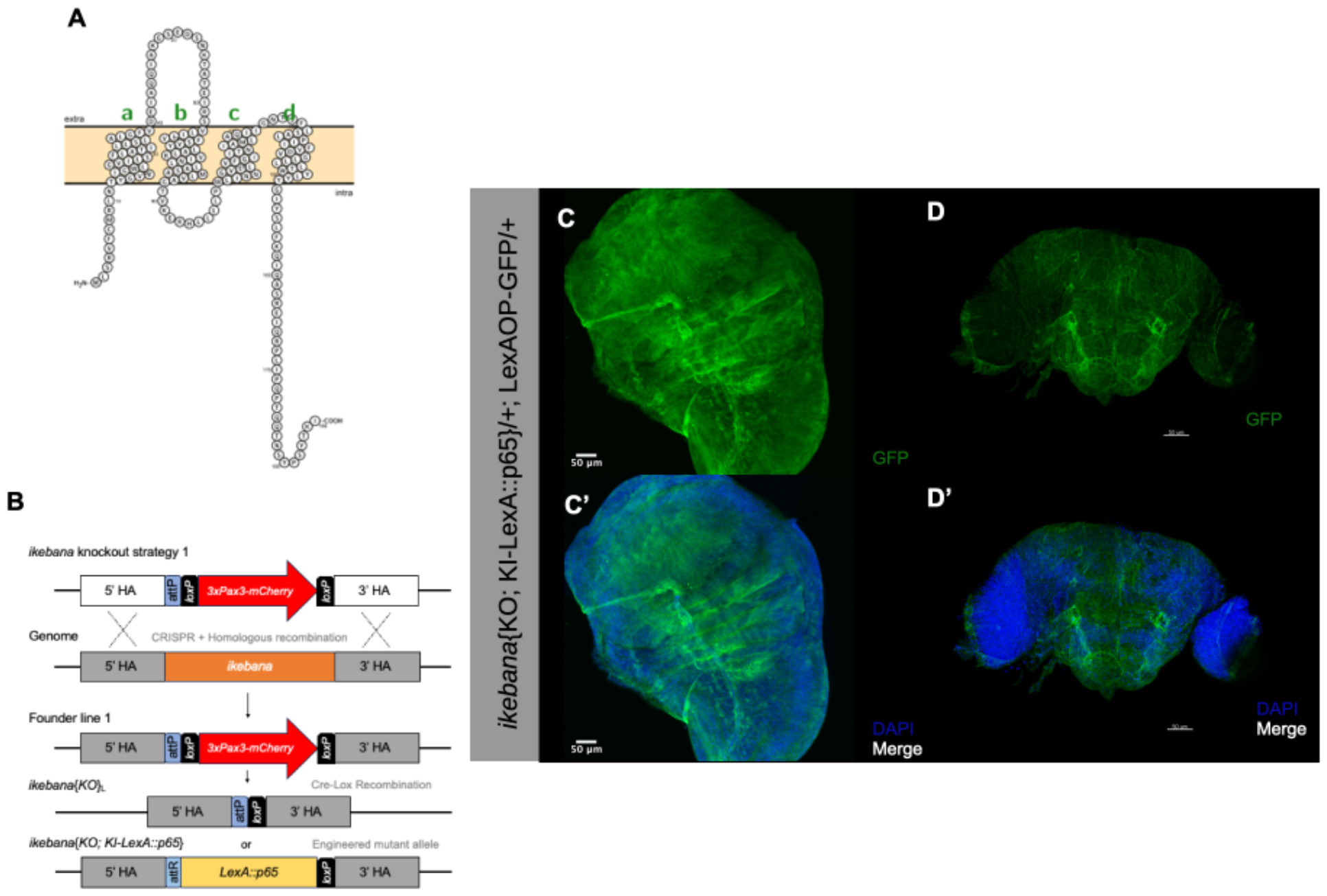
Ikebana is basally expressed in several cells of the wing imaginal disc and adult brain. (A) Predicted configuration of the Ikebana protein at the cell membrane. The green letters a-d indicate the four transmembrane domains represented by the green letters a-d. Model done with Protter. (B) Strategy for the generation of the *ikebana*{KO}L and, consequently, the *ikebana*{KO; KI-LexA::p65} transgenic lines. The entire *ikebana* coding sequence in the genome is removed by CRISPR with two sgRNA (Baena-Lopez et al. 2013). Then the genome is repaired, by homologous recombination, allowing the introduction of the *3xPax3-mCherry* selection marker (Founder line 1) (Huang et al. 2009). For the *ikebana*{KO}L transgenic line, the *3xPax3-mCherry* selection marker was removed by Cre-lox recombination. For the generation of the *ikebana*{KO; KI-LexA::p65} transgenic line, a knockin construct containing the LexA::p65 sequence under the control of the endogenous *ikebana* promoter, was integrated into the *ikebana* knockout locus. The pTV3 vector backbone was removed in the final knockin line. (C-D) Expression of Ikebana in the wing imaginal discs of third-instar larvae (C) and the adult brain (D). GFP, in green, marks the cells trying to express ikebana, and DAPI, in blue, marks the cell nuclei. Scale bars, 50μm. Genotype: *ywF*; *ikebana*{KO; KI-*LexA::p65*}/+; *26xLexAop-CD8::GFP*/+.

To evaluate the expression of *ikebana*, we employed CRISPR-Cas9 technology, coupled with homologous recombination to generate a genetic *ikebana* knockout (KO) line and introduce a genetic marker to allow its identification (Baena-Lopez et al. 2013; Huang et al. 2009). In this *ikebana* KO line, we inserted a LexA::p65 sequence under the control of the *ikebana* endogenous promoter, which, in conjunction with LexAop-GFP, facilitated the specific labelling of cells trying to express *ikebana* (*Figure* 1B). We found that *ikebana* exhibits endogenous widespread expression across third-instar larvae wing imaginal discs and within the adult brain (*Figure* 1C).

### Ikebana partially protects loser cells from elimination

To elucidate the role of Ikebana in cell competition, we conducted experiments to investigate the consequences of modulating the expression of *ikebana* in loser clones. For this purpose, we used the supercompetition assay, which relies on the *tub>dmyc>Gal4* cassette (Moreno and Basler 2004). Upon activating a heat shock Flippase, this system facilitates the recombination of FRT sites, allowing the expression of both GFP and a construct of interest. By manipulating the heat shock duration, we generated GFP-marked clones scattered within cells carrying an additional copy of *dmyc*. This supplementary *myc* copy designates the background cells as winners, while the clones lacking this additional copy are losers whose elimination can be prevented by downregulating the *flower lose* isoforms or overexpressing *diap1* (Death-associated inhibitor of apoptosis 1).

Contrary to downregulating *flower loseA/B*, downregulating *ikebana* in the loser clones did not block their elimination at 72 hours after clone induction (ACI), as shown in *Figure* 2A-D. However, the overexpression of *ikebana* in the loser clones significantly reduced their elimination compared to the negative control at 72 hours ACI (*Figure* 2E-H). Notably, this protective effect was not as pronounced as when overexpressing *diap1*, suggesting that Ikebana confers partial protection to loser cells from elimination.

**Figure 2.**
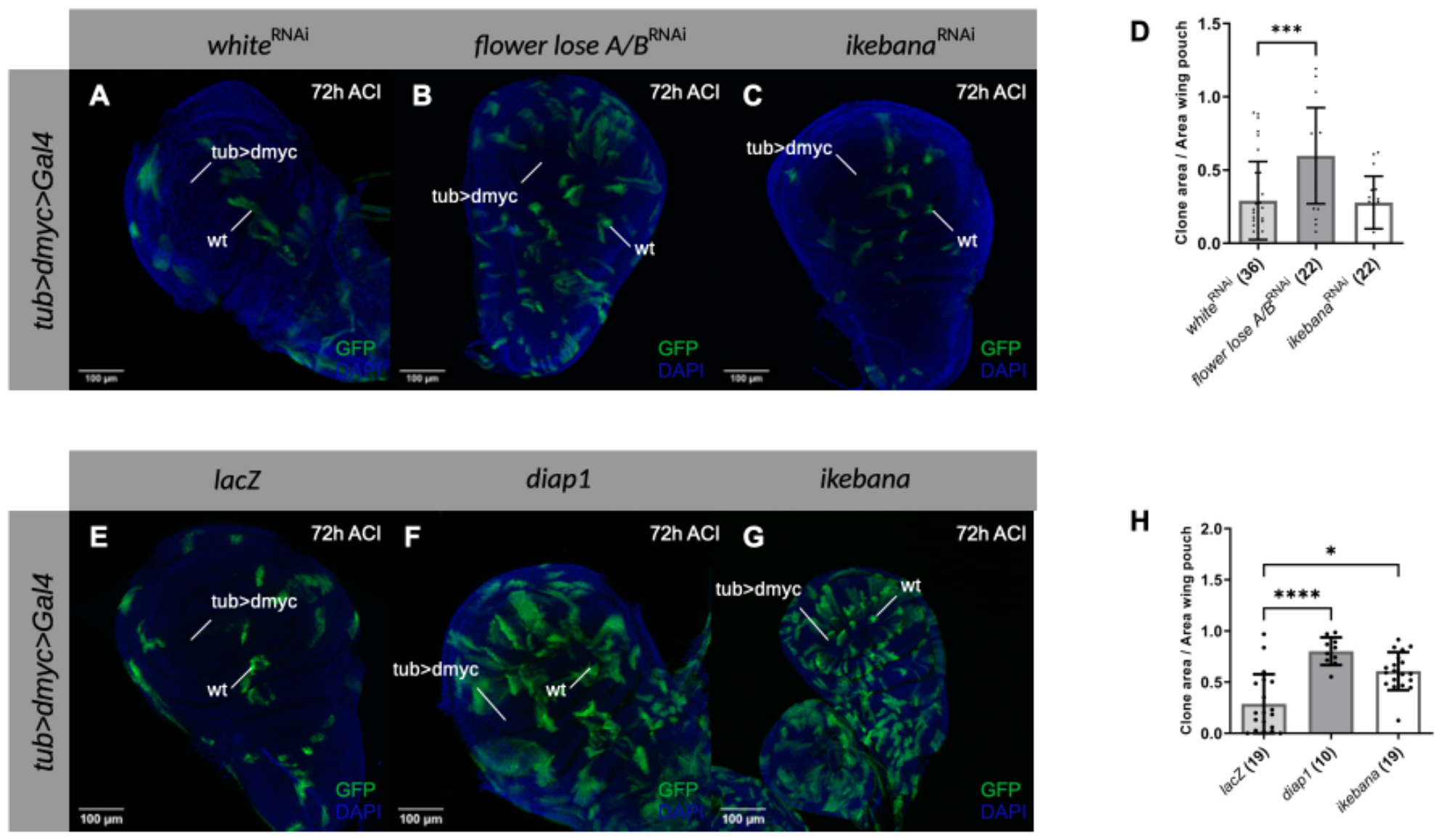
Ikebana partially protects loser cells from elimination. (A-C) Supercompetition assay in the wing imaginal discs of third instar larvae in which loser wild-type clones (GFP) are outcompeted by *dmyc* overexpressing cells (Rhiner et al. 2010). UAS-*white-*RNAi (A), UAS-*flower lose A/B-*RNAi (B) and UAS-*ikebana-*RNAi (C) are expressed in the loser clones 72h after clone induction (ACI). DAPI is represented in blue. Scale bars, 100 μm. (D) Quantification of the clone area over the total area of the wing pouch when UAS-*white-*RNAi, UAS-*flower lose A/B-*RNAi, and UAS-*ikebana-*RNAi are expressed in the loser clones 72h ACI. We normalised the ratio at 12 hours ACI (data not shown), and subsequently, the 72h time point is normalised relative to their respective conditions at 12 hours. (E-G) Supercompetition assay in the wing imaginal discs of third instar larvae in which loser wild-type clones (GFP) are outcompeted by *dmyc* overexpressing cells (Rhiner et al. 2010). UAS-*lacZ* (A), UAS-*diap1* (B) and UAS-*ikebana* (C) are expressed in the loser clones 72h ACI. DAPI is represented in blue. Scale bars, 100 μm. (H) Quantification of the clone area over the total area of the wing pouch when UAS*-lacZ*, UAS-*diap1*, and UAS-*ikebana* are expressed in the loser clones 72h ACI. We normalised the ratio at 24 hours ACI (data not shown), and subsequently, the 72h time point is normalised relative to their respective conditions at 24 hours. The numbers after the genotypes indicate the number of discs analysed. Error bars indicate SD; ^*^P<0.05; ^***^P<0.001; ^****^P<0.0001. Statistical significance between groups was calculated using the nonparametric Kruskal-Wallis test and a Dunn’s test was applied for multiple comparisons between genotypes. Genotypes: *ywF/w*; *tub>dmyc>Gal4*, UAS-*GFP*/+; UAS-*white*-RNAi/MKRS (A); *ywF/w*; *tub>dmyc>Gal4*, UAS-*GFP*/+; UAS-*flowerloseA/B*-RNAi/MKRS (B); *ywF/w*; *tub>dmyc>Gal4*, UAS-*GFP*/UAS-*ikebana*-RNAi; MKRS/+ (C); *ywF/w*; *tub>dmyc>Gal4*, UAS-*GFP*/UAS-*lacZ*; MKRS/+ (E); *ywF/w*; *tub>dmyc>Gal4*, UAS-*GFP*/+; UAS-*diap1*/MKRS (F); *ywF/w*; *tub>dmyc>Gal4*, UAS-*GFP*/+; UAS-*ikebana*/MKRS (G);

To ascertain that this effect is specific to cell competition, we conducted experiments to exclude the possibility that Ikebana is a general apoptosis regulator. First, we expressed *eiger* – cell death initiator via JNK signalling in the eye, which resulted in a reduced eye phenotype (Igaki et al. 2002). We then examined whether Ikebana could rescue the normal phenotype (Figure S*1*A-L). Additionally, considering that the rotation of the fly terminalia during development is apoptosis-dependent (Benitez et al. 2010), we investigated whether overexpressing or downregulating *ikebana* in this region would impact normal terminalia rotation, manifesting as incomplete terminalia rotation in the adult fly (Figure S*1*M-S). Results show that modulating *ikebana* expression does not interfere with the eye phenotype or terminalia rotation (Figure S*1*), indicating that Ikebana is not a general regulator of apoptosis, and that the effects observed in clones are indeed attributable to cell competition.

### Ikebana regulates Flower LoseB expression

Given the basal expression of *ikebana* in the wing imaginal discs of the third instar larvae, we explored whether downregulating its expression with *ikebana*-RNAi can induce cell competition. Using the *act>y+>Gal4* cassette, we generated GFP-labeled wild-type clones surrounded by cells expressing an additional copy of *yellow*, which are also wild-type in the context of cell competition. As a negative control, we expressed *white*-RNAi in the clones, which is not predicted to interfere with cell competition, and, as a positive control, we overexpressed *flower loseB*, which will confer a loser state to the clones, resulting in their elimination over time.

At 72 hours ACI, *ikebana*-RNAi expression resulted in a reduced clonal area compared to the *white*-RNAi control. Such reduction of the clonal area was even more pronounced than the one observed with the overexpression of *flower loseB* (*Figure* 3A-D). This suggests that decreasing *ikebana* expression is sufficient to induce a loser state, although more experiments are required to prove that this is due to cell competition.

**Figure 3.**
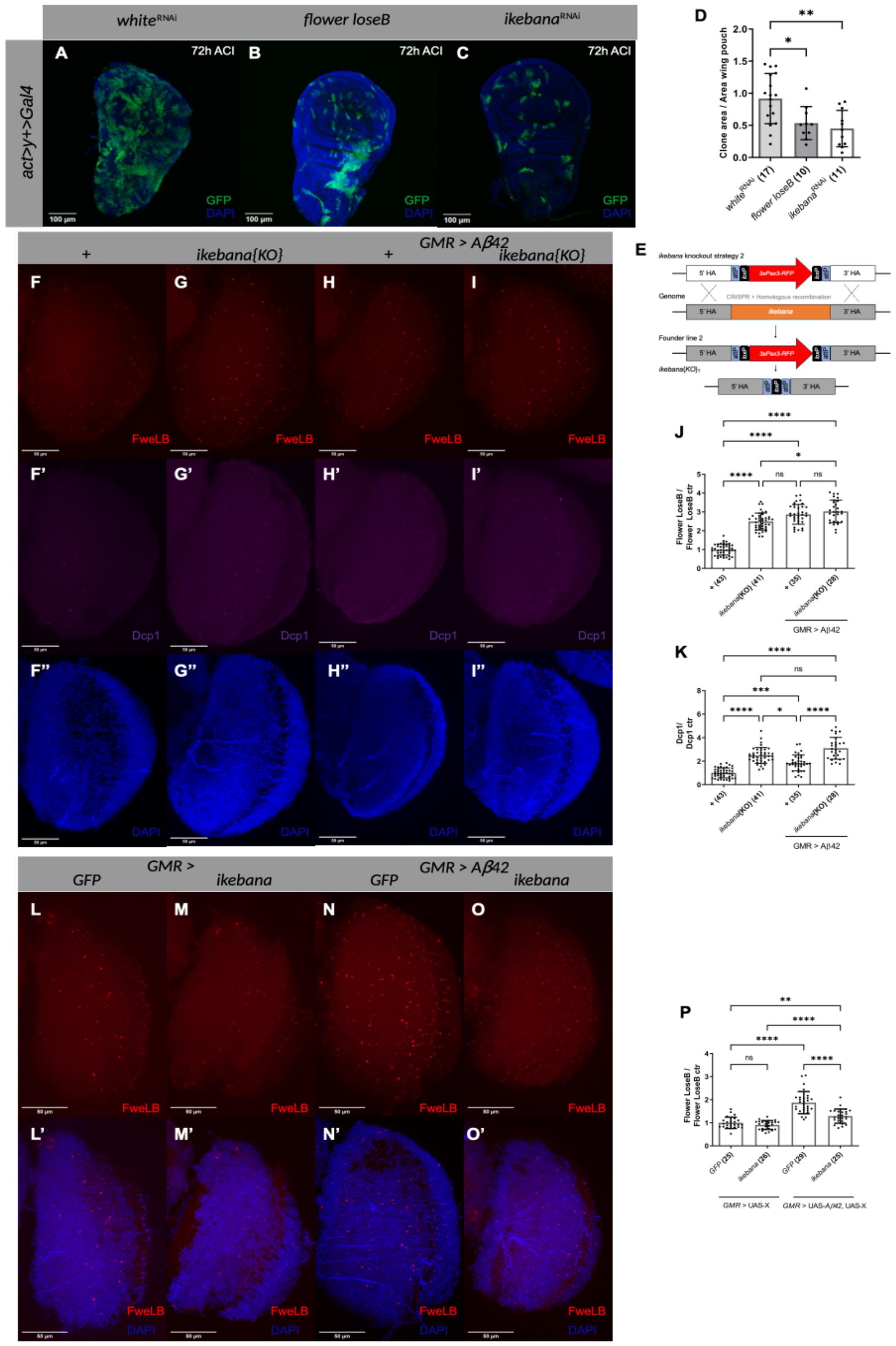
Ikebana modulates Flower LoseB expression. (A) Competition assay in the wing imaginal discs of third instar larvae in which wild-type clones (GFP) are in a background of *yellow* expressing cells (Rhiner et al. 2010). UAS-*white-*RNAi (A), UAS-flowerloseB (B) and UAS-*ikebana-*RNAi (C) are expressed in the wild-type clones 72h ACI. DAPI is represented in blue. Scale bars, 100 μm. Genotypes: *ywF*; *act>y>Gal4*, UAS-*GFP*/+; UAS-*white*-RNAi/MKRS (A); *ywF*; *act>y>Gal4*, UAS-*GFP*/UAS-*flowerloseB*; MKRS/+ (B); *ywF*; *act>y>Gal4*, UAS-*GFP*/UAS-*ikebana-*RNAi; MKRS/+ (C) (D) Quantification of the clone area over the total area of the wing pouch UAS-*white-*RNAi, UAS-flowerloseB and UAS-*ikebana-*RNAi are expressed in the loser clones 72h ACI. We normalised the ratio at 24 hours ACI (data not shown), and subsequently, the 72h time point is normalised relative to their respective conditions at 24 hours. (E) Strategy for the generation of the *ikebana*{KO}T transgenic line. The entire *ikebana* coding sequence in the genome is removed by CRISPR with two sgRNA (Baena-Lopez et al. 2013). Then the genome is repaired, by homologous recombination, allowing the introduction of the *3xPax3-RFP* selection marker (Founder line 2) (Huang et al. 2009). The *3xPax3-RFP* marker was removed by Cre-lox recombination. (F-I) Expression of Flower LoseB and Dcp1 in *ikebana* KO adult optic lobes. Optic lobe of control flies (F); *ikebana*{KO} (G); *GMR>Aβ42* (H) and *GMR>Aβ42, ikebana*{KO} (I). Flower LoseB is marked in red; α-Dcp1 is marked in magenta; DAPI, in blue, marks the cell nuclei. 41-μm image projections. Scale bars, 50 μm. Genotypes: *w*; ; *flower*{KO; KI-*flowerloseB::mCherry*}/+ (F); *w*; *ikebana*{KO}T/*ikebana*{KO}L; *flower*{KO; KI-*flowerloseB::mCherry*}/+ (G); *w*; *GMR-Gal4, UAS-Aβ42*/+; *flower*{KO; KI-*flowerloseB::mCherry*}/+ (H); *w*; *GMR-Gal4, UAS-Aβ42, ikebana*{KO}T/*ikebana*{KO}L; *flower*{KO; KI-*flowerloseB::mCherry*}/+ (I). (J) Quantification of the number of Flower LoseB positive cells in the optic lobes normalised against the control (ctr) +. (K) Quantification of the number of Dcp1 positive cells in the optic lobes normalised against the ctr +. (L-O) Expression of Flower LoseB in the adult optic lobes when *ikebana* is overexpressed. Optic lobe of control flies overexpressing *GFP* (L) or *ikebana* (M), or flies expressing the Aβ42 peptide in the GMR region, also overexpressing *GFP* (N) or *ikebana* (O). Flower LoseB is marked in red; DAPI, in blue, marks the cell nuclei. 41-μm image projections. Scale bars, 50 μm. Genotypes: *w*; *GMR-Gal4*/*UAS-CD8::GFP*; *flower*{KO; KI-*flowerloseB::mCherry*}/+ (L); *w*; *GMR-Gal4*/+; *flower*{KO; KI-*flowerloseB::mCherry*}/UAS-*ikebana* (M); *w*; *GMR-Gal4*, UAS-A*β42*/*UAS-CD8::GFP*; *flower*{KO; KI-*flowerloseB::mCherry*}/+ (N); *w*; *GMR-Gal4, UAS-Aβ42*/+; *flower*{KO; KI-*flowerloseB::mCherry*}/UAS-*ikebana* (O). (P) Quantification of the number of Flower LoseB positive cells normalised against the control GFP in the ctr scenario. The numbers after the genotypes indicate the number of discs or optic lobes analysed. Error bars indicate SD; ns indicates non-significant; ^*^P<0.05; ^**^P<0.01; ^***^P<0.001; ^****^P<0.0001. Statistical significance between groups was calculated using the nonparametric Kruskal-Wallis test and a Dunn’s test was applied for multiple comparisons between genotypes (D, J and K). Statistical significance between groups was calculated using the Brown-Forsythe test and a Welch ANOVA test and Dunnette’s T3 was applied for multiple comparisons between genotypes (P).

We then asked whether Ikebana regulates the loser state of a cell by influencing Flower expression. Using a FlowerLoseB::mCherry reporter line, we assessed its production in an *ikebana* KO scenario (Figure 3E-K). For the *ikebana* KO condition, we crossed the *ikebana* KO_T_ with the *ikebana* KO_L_, a transheterozygous allelic version, because we observed the presence of off-target effects in the *ikebana* KO_L_ line (data not shown). In the optic lobe of *ikebana* KO flies, we found a 2.5-fold increase in Flower LoseB-positive cells compared to the control wild type for *ikebana* (*Figure* 3J). This rise in the number of loser cells was accompanied by increased Dcp1-positive cells (*Figure* 3K), meaning these cells were actively being eliminated.

We then tested an Alzheimer’s disease model scenario where we expected more loser cells (Coelho et al. 2018). Intriguingly, in this scenario, removing *ikebana* did not further increase the number of loser cells in the optic lobe (*Figure* 3J). However, it did increase the number of dying cells (*Figure* 3K), as measured via Dcp-1 positivity. We also examined whether the same happened for Azot expression, and indeed, the absence of *ikebana* is sufficient to increase the number of Azot-positive cells in basal cell competition but not in an Alzheimer’s disease model (Figure S*2*). These data suggest that the absence of *ikebana* alone is often sufficient to increase the number of loser cells. However, in an Alzheimer’s disease model, Ikebana prevents cell elimination without affecting Flower LoseB expression, as seen by the increase in dying cells in the *ikebana* KO scenario without an increase in the number of loser cells.

Lastly, we investigated whether overexpressing *ikebana* in the optic lobe, using the *GMR* driver, would decrease the number of Flower LoseB-positive cells (*Figure* 3L-P). Under basal levels of cell competition, overexpressing *ikebana* did not affect the number of Flower LoseB-positive cells. However, in an Alzheimer’s disease model, overexpressing *ikebana* in the *GMR* region led to a 0.7-fold decrease in the number of cells producing Flower LoseB::mCherry compared to the GFP control (*Figure* 3P). This suggests that high levels of Ikebana can reverse the low fitness status of cells, indicating that Ikebana is sufficient to partially block the loser fate specification in the Alzheimer’s disease model.

## Discussion

Ikebana, anticipated as a transmembrane protein which physically interacts with Flower, emerges as a pivotal contributor to cell competition dynamics. Using a clonal assay, we observed that localised reduction of *ikebana* expression within clones results in a diminished clonal area, similar to local overexpression of *flower loseB*, which forces the cells into a loser state. Given the seemingly ubiquitous expression of *ikebana* in wing imaginal discs, we propose that, for a cell to adopt a loser state, it must first downregulate *ikebana*. We hypothesise that this decrease in Ikebana production will promote the production of Flower LoseB, leading to cell elimination. In adult brains, the absence of *ikebana* proves sufficient to increase the number of cells positive for Flower LoseB or Azot, further supporting this hypothesis. In scenarios anticipating a high number of loser cells, as in the Alzheimer’s disease model, overexpressing *ikebana* correlates with a decrease in the number of Flower LoseB-positive cells, implying that Ikebana can revert the low fitness of the cells - a refined mechanism which likely operates to selectively eliminate the necessary number of losers, preserving tissue function. In such cases, Ikebana can be re-expressed in loser cells, reverting their loser state by decreasing Flower LoseB expression and sparing selected loser cells from elimination (*Figure* 4).

**Figure 4.**
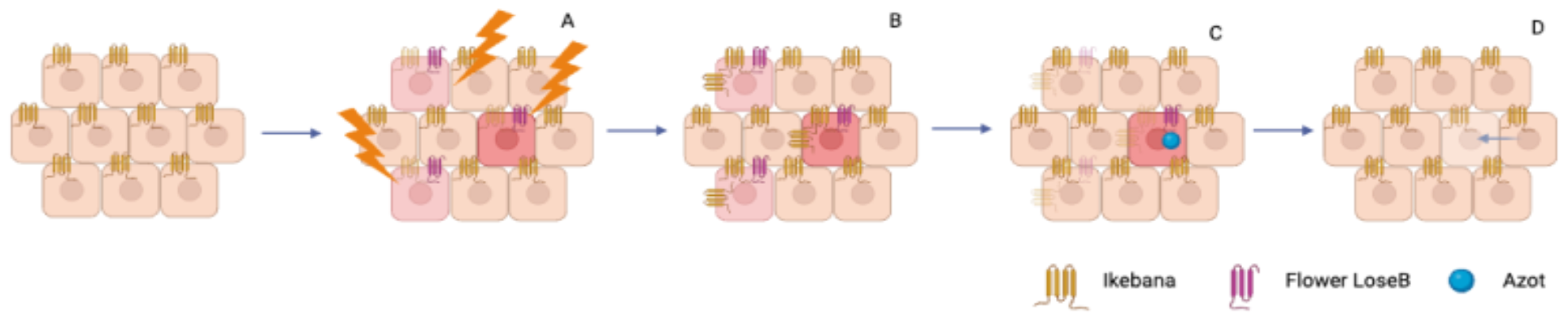
Model of the action of Ikebana in cell competition. When an insult occurs, *ikebana* is downregulated, causing the suboptimal cells to express Flower LoseB and be labelled as loser (A). The loser cells that are not supposed to be eliminated express *ikebana* again and revert their loser status (B). The loser cells that should be eliminated express Azot and activate the apoptotic machinery (C). The loser cells are eliminated and, in some cases, replaced by fitter cells (D). Model done with Biorender.com.

To strengthen the model, additional experiments are needed to exclude interference with cell proliferation. Overexpressing or downregulating *ikebana* in specific regions and comparing sizes against a control without *ikebana* manipulation, coupled with immunostaining for cell death in clones, would bolster the robustness of our conclusions. Although these experiments were not conducted, observational evidence using the *actin-Gal4* driver indicates normal development time and adult fly size when manipulating *ikebana* (data not shown). This supports our conviction that the observed effects are due to cell competition rather than mere differences in proliferation rates.

Ikebana joins a growing list of proteins, such as SPARC, implicated in protecting loser cells during Flower-dependent cell competition (Portela et al. 2010). However, SPARC does not belong to the Flower-dependent cell competition canonical pathway because manipulating *flower* in loser or wild-type clones does not change the levels of SPARC (Portela et al. 2010). On the contrary, Ikebana is proposed to interact directly with Flower. Additionally, *SPARC* is exclusively expressed in loser cells to prevent their elimination, whereas *ikebana* appears ubiquitously expressed.

Interestingly, Ikebana was previously identified as a member of the tetraspanins-pasiflora family of proteins, characterised by four transmembrane domains and conserved regions (Deligiannaki et al. 2015). In *Drosophila*, only two other proteins, Pasiflora and Fire Exit, are characterised, albeit in embryos, and their role in cell competition remains unexplored (Deligiannaki et al. 2015; With et al. 2003). This protein family extends to humans, including the lysosomal-associated proteins LAPTM4A and LAPTM4B. Ikebana shares more structural similarities with LAPTM4A (Deligiannaki et al. 2015). This human protein, associated with the clearance of proteins from the plasma membrane via endocytosis and implicated in inducing multidrug resistance in *Saccharomyces cerevisiae*, presents a promising avenue for therapeutic exploration (Grabner et al. 2011; Hogue, Kerby, and Ling 1999). Future work is needed to confirm LAPTM4A as the human orthologue of Ikebana and test if it is involved in the clearance of Flower lose isoforms from the cell membrane. If so, manipulation of LAPTM4A could open therapeutic avenues to prevent the unnecessary elimination of loser cells.

## Experimental Procedures

### Drosophila Genetics and experimental setups

Stocks and crosses were kept at 25ºC in Vienna standard media with extra yeast. All stocks were obtained from Bloomington Stock Center unless specified. The following RNAi lines from VDRC were used: UAS-*ikebana*-RNAi (ID 111608, Chr2, viable), UAS-*eiger*-RNAi (ID 12658, Chr3, viable), UAS-*azot*-RNAi (ID 7219, Ch3, viable). For the overexpression of *ikebana*, a UAS-*ikebana* stock was generated according to standard procedures.

For the supercompetition assay, the following stocks were used: *tub>dmyc>Gal4*, UAS-*GFP*, UAS-*white*-RNAi, UAS-*flowerloseA/B-*RNAi, UAS-*lacZ*, and UAS-diap1. The larvae were given a 17-minute heat shock at 37ºC, and the vials were placed at 29ºC until dissected.

For the wild-type clones in the wild-type background, the following stocks were used: *act>y+>Gal4*, UAS-*GFP, and UAS-flowerloseB*. The larvae were given an 8-minute heat shock at 37ºC, and the vials were placed at 29ºC until dissected.

For the experiment to understand the effects of *ikebana* KO in the number of Flower LoseB or Azot-positive cells, the following stocks were used: *white*; *ikebana{KO}T/ikebana{KO}L*; *GMR-Gal4, UAS-Aβ42*; *flower{KO;KI-flowerloseB::mCherry*}; *azot::mCherry*. After hatching, female flies were kept at 29ºC until 2 weeks old, when they were dissected.

For the experiment to understand the effects of overexpressing *ikebana* in the number of Flower LoseB-positive cells, the following stocks were used: *GMR-Gal4* and *UAS-GFP*. After hatching, female flies were kept at 29ºC until 2 weeks old, when they were dissected.

For the experiments to understand if Ikebana is a general regulator of apoptosis, the following stocks were used: *GMR-Gal4, UAS-eiger* and *Engrailed-Gal4*. The eyes or genitalia were observed in the first week of adulthood.

### Ikebana knockout-L Generation

CRISPR-mediated mutagenesis followed by homologous recombination was performed by the Fly Platform at the Champalimaud Foundation, using the methods highlighted in Baena-Lopez et al. 2013. In brief, the upstream gRNA sequence ACCAACTGCTTGAACCA[GTC] and the downstream gRNA sequence [GCT]ACACCAAGATTTAAGCT were cloned into a pCFD5 vector. The cassette 3xPax3::mCherry contains one attP site, a floxed 3xPax3::mCherry, and two homology arms were cloned into pTV3 as a donor template for repair.

CG15098-targeting gRNAs and a donor plasmid were microinjected into embryos nos-Cas9. F1 flies carrying the selection marker 3xPax3::mCherry were further validated by genomic PCR and sequenced. CRISPR generates an 1139-bp deletion allele of CG15098, deleting the partial 5’ UTR/3’ UTR and entire CDS of the CG15098 gene and replacing it with cassette 3xPax3::mCherry. This cassette was then removed by Cre-lox recombination, leaving only the attP site and the loxP.

### Ikebana knockout-T Generation

CRISPR-mediated mutagenesis was performed by WellGenetics Inc. using modified methods of Kondo and Ueda 2013. Briefly, the upstream gRNA sequence GTGAATCCAGAATGCTGTCC[AGG] and the downstream gRNA sequence GGCCAAACGGGAAGCTACAC[TGG] were cloned into U6 promoter plasmid(s) separately. Cassette RMCE-3xPax3::RFP contains two attP sites, a floxed 3xPax3::RFP, and two homology arms were cloned into pUC57-Kan as donor templates for repair.

CG15098-targeting gRNAs and hs-Cas9 were supplied in DNA plasmids and a donor plasmid for microinjection into embryos of control strain *w*[1118]. F1 flies carrying the selection marker 3xPax3::RFP were further validated by genomic PCR and sequencing. CRISPR generates a 1029-bp deletion allele of CG15098, deleting partial 5’ UTR/3’ UTR and entire CDS of CG15098 gene and is replaced by cassette RMCE-3xPax3::RFP. This cassette was then removed by Cre-lox recombination, leaving only the attP sites and the loxP.

### Ikebana knockin Generation

For the generation of the *ikebana*{KO; KI-*LexA::p65*}, the cDNA of LexA::p65 was generated and inserted into a RIV^Cherry^ vector (Baena-Lopez et al. 2013), and the knockin was generated as described in Huang et al. 2009. Primer sequences are available upon request.

### Immunohistochemistry and image acquisition

Wing imaginal discs of third-instar larvae were dissected in chilled PBS, fixed for 20 minutes in formaldehyde (4% v/v in PBS), and permeabilised with PBT 0.4% Triton.

Adult brains were dissected in chilled PBS; the samples were fixated for 30 minutes in formaldehyde (4% v/v in PBS) and permeabilised with PBT 1% Triton. The wing imaginal discs or brains were then incubated with rabbit α-Dcp1 (1:50) from Cell Signaling (#9578). Samples were mounted in Vectashield with DAPI (Vectorlabs).

Confocal images were acquired with Zeiss LSM 880 using the Plan-Apochromat 20x/0.8 M27 dry objective for the case of the wing imaginal discs and the Plan-Apochromat 40x/1.4 Oil DIC M27 objective for the case of the adult brains. Maximum intensity projections of the wing imaginal discs or 41-μm-wide images of the adult brains were obtained with Zeiss Zen Black.

To capture images of the adult eyes and genitalia, the Leica S9 I Stereomicroscope was used. Flies were either anaesthetised or imaged alive in a CO_2_ station.

### Quantification and statistical analysis

Fiji-ImageJ macros developed in this work, available upon request, quantified the clone areas as the number of Flower LoseB, Azot, or Dcp1-positive cells. The areas of the adult eyes were measured in Fiji-ImageJ. For each condition, a minimum of 10 wing imaginal discs, 23 optic lobes, and 12 adult eyes were analysed.

Statistical significance between groups was calculated using the nonparametric Kruskal-Wallis test, and Dunn’s test was applied for multiple comparisons between genotypes. Specifically in Figure 3P, statistical significance between groups was calculated using the Brown-Forsythe and Welch ANOVA tests, and Dunnett’s T3 was applied for multiple comparisons between genotypes.

## Abbreviations

Aβ42: Amyloid beta 42
ACI: After Clone Induction
bp: base pair
ctr: control
Diap1: Death-associated inhibitor of apoptosis 1
KI: knockin
KO: knockout
LAPTM: Lysosomal Protein Transmembrane

## Author Contributions

M.M.R., A.G.G., and E.M. designed the experiments. M.M.R. performed and analysed the experiments. A.G.G. helped with image acquisition and statistical analysis. C.B.P. helped with data analysis and design of the transgenic constructs. C.F.C., I.A. and M.S.E obtained preliminary data. B.H. developed the UAS-*ikebana* transgenic flies. M.M.R. wrote the manuscript.

## Acknowledgements

We thank Bloomington Stock Center for flies; WellGenetics Inc. for the generation of the *ikebana* KO_T_; the technicians at the Champalimaud Fly Platform for support with stock maintenance; Catarina Craveiro and the MTT platform for support in the generation of transgenic flies, the ABBE platform for microscopy support and Andrea Spinazzola for his suggestions and comments on the manuscript. M.R was supported by an FCT - Fundação para a Ciência e a Tecnologia - PhD studentship (SFRH/BD/138537/2018). This study was supported by Portuguese national funds, through FCT in the context of the project UIDB/04443/2020 and the European Research Council (Consolidator Grant to E.M.: ‘‘Active Mechanisms of Cell Selection: From Cell Competition to Cell Fitness’’). Fly and MTT platforms were funded by the research infrastructure CONGENTO, co-financed by Lisboa Regional Operational Programme (Lisboa2020), under the PORTUGAL 2020 Partnership Agreement, through the European Regional Development Fund (ERDF) and Fundação para a Ciência e Tecnologia (Portugal) under the project LISBOA-01-0145-FEDER-022170. The Portuguese Platform of Bioimaging funded ABBE platform - LISBOA-01-0145-FEDER-022122.

## Figures

**Figure S1.**
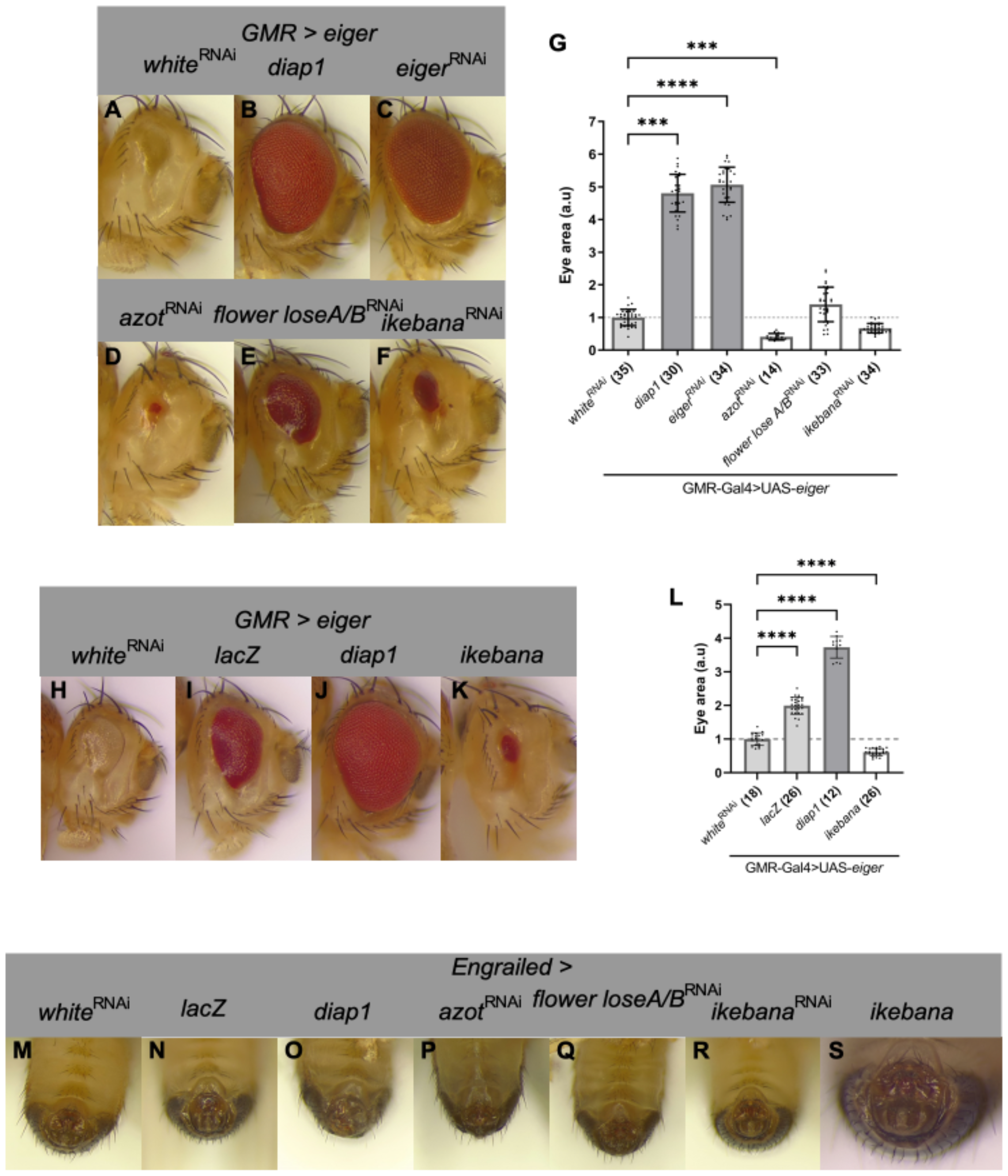
Ikebana is not a general regulator of apoptosis. (A-F) Representative eyes of *Drosophila* when UAS-*white*-RNAi (A), UAS-*diap1* (B), UAS-*eiger*-RNAi (C), UAS-*azot*-RNAi (D), UAS-*flowerloseA/B*-RNAi (E) and UAS-*ikebana-*RNAi (F) are co-expressed with Eiger in the GMR domain. Genotypes: *yw/w*; *GMR-Gal4, UAS-eiger*/+; UAS-*white*-RNAi/+ (A); *yw/w*; GMR-Gal4, UAS-*eiger*/+; UAS-*diap1*/+ (B); *yw/w*; *GMR-Gal4*, UAS-*eiger*/+; UAS-*eiger*-RNAi/+ (C) *yw/w*; *GMR-Gal4*, UAS-*eiger*/+; UAS-*azot*-RNAi/+ (D); *yw/w*; *GMR-Gal4*, UAS-*eiger*/+; UAS-*flowerloseA/B*-RNAi/+; (E) *yw/w*; *GMR-Gal4*, UAS-*eiger*/UAS-*ikebana*-RNAi; +/+ (F). (G) Quantification of the eye area of the flies (A-F) normalised for *white*-RNAi. (H-K) Representative eyes of Drosophila when UAS-*white*-RNAi (H), UAS-*lacZ* (I), UAS-*diap1* (J) and UAS-*ikebana* (K) are co-expressed with Eiger in the GMR domain. Genotypes: *yw/w*; *GMR-Gal4, UAS-eiger*/+; UAS-*white*-RNAi/+ (H); *yw/w*; *GMR-Gal4, UAS-eiger*/UAS-lacZ; +/+ (I); *yw/w*; *GMR-Gal4, UAS-eiger*/+; UAS-diap1/+ (J); *yw/w*; *GMR-Gal4, UAS-eiger*/+; UAS-*ikebana*/+ (K); (K) Quantification of the eye area of the flies (H-K) normalised for *white*-RNAi. Horizontal line when eye area = 1 arbitrary unit (a.u) to show the differences of the other genotypes against the control. The numbers after the genotypes indicate the number of eyes analysed. Error bars indicate SD; ^***^P<0.001; ^****^P<0.0001. Statistical significance between groups was calculated using the nonparametric Kruskal-Wallis test and a Dunn’s test was applied for multiple comparisons between genotypes. (M-S) Representative images of the genitalia of the fly when UAS-*white*-RNAi (M), UAS-*lacZ* (N), UAS-*diap1* (O), UAS-*azot*-RNAi (P), UAS-*flowerloseA/B*-RNAi (Q), UAS-*ikebana*-RNAi (R) and UAS-*ikebana* (S) are expressed under the control of the *engrailed* driver Genotypes: *ywF/w*; *engrailed-Gal4, UAS-GFP*/+; UAS-*white*-RNAi/+ (M); *ywF/w*; *engrailed-Gal4, UAS-GFP*/UAS-*lacZ*; +/+ (N); *ywF/w*; *engrailed-Gal4, UAS-GFP*/+; UAS-*diap1*/+ (O); *ywF/w*; *engrailed-Gal4, UAS-GFP*/+; UAS-*azot*-RNAi/+ (P); *ywF/w*; *engrailed-Gal4, UAS-GFP*/+; UAS-*flowerloseA/B*-RNAi/+ (Q); *ywF/w*; *engrailed-Gal4, UAS-GFP*/UAS-*ikebana*-RNAi; +/+ (R); *ywF/w*; *engrailed-Gal4, UAS-GFP*/+; UAS-*ikebana*/+ (S);

**Figure S2.**
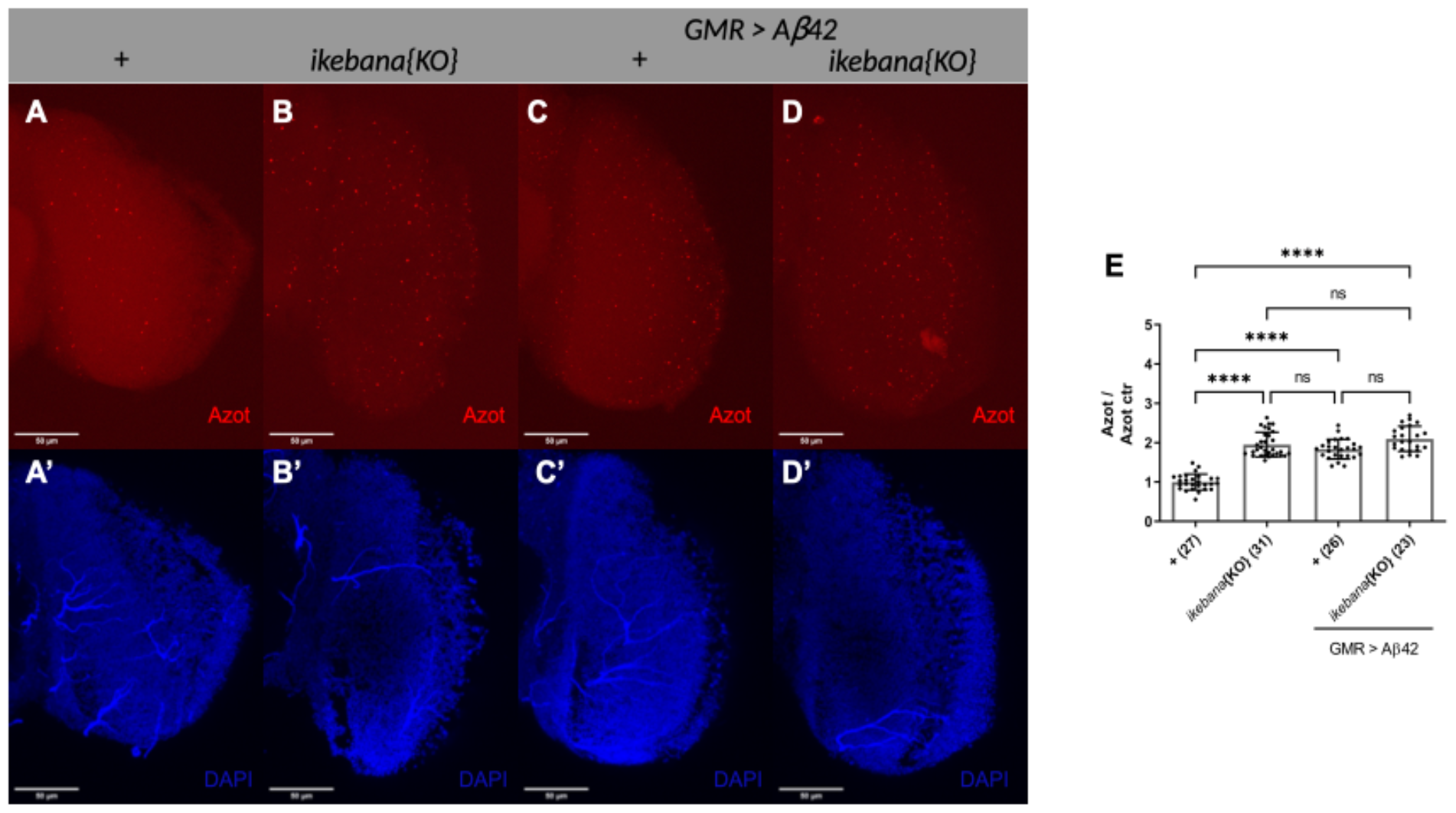
Ikebana modulates Azot expression. (A-D) Expression of Azot in *ikebana* KO adult optic lobes. Optic lobe of control flies (A); *ikebana*{KO} (B); *GMR>Aβ42* (C) and *GMR>Aβ42, ikebana*{KO} (D). Azot is marked in red; DAPI, in blue, marks the cell nuclei. 41-μm image projections. Scale bars, 50 μm. Genotypes: *w*; ; *azot::mCherry*/+ (A); *w*; *ikebana*{KO}T/*ikebana*{KO}L; *azot::mCherry*/+ (B); *w*; *GMR-Gal4, UAS-Aβ42*/+; *azot::mCherry*/+ (C); *w*; *GMR-Gal4, UAS-Ab42, ikebana*{KO}T/*ikebana*{KO}L; *azot::mCherry*/+ (D). (E) Quantification of the number of Azot positive cells in the optic lobes normalised against the ctr +. The numbers after the genotypes indicate the number of discs or optic lobes analysed. Error bars indicate SD; ns indicates non-significant; ^****^P<0.0001. Statistical significance between groups was calculated using the nonparametric Kruskal-Wallis test and a Dunn’s test was applied for multiple comparisons between genotypes.

## Notes

### Competing Interest Statement

The authors have declared no competing interest.

